# SARS-CoV-2 infection, disease and transmission in domestic cats

**DOI:** 10.1101/2020.08.04.235002

**Authors:** Natasha N. Gaudreault, Jessie D. Trujillo, Mariano Carossino, David A. Meekins, Igor Morozov, Daniel W. Madden, Sabarish V. Indran, Dashzeveg Bold, Velmurugan Balaraman, Taeyong Kwon, Bianca Libanori Artiaga, Konner Cool, Adolfo García-Sastre, Wenjun Ma, William C. Wilson, Jamie Henningson, Udeni B. R. Balasuriya, Juergen A. Richt

**Author notes:** Department of Veterinary Pathobiology and Department of Molecular Microbiology and Immunology, University of Missouri, Columbia, MO, USA. Corresponding author: Dr. Juergen A. Richt; Department of Diagnostic Medicine/Pathobiology, College of Veterinary Medicine, Kansas State University, Manhattan, KS, USA.

## Abstract

Severe Acute Respiratory Syndrome Coronavirus 2 (SARS-CoV-2) is the cause of Coronavirus Disease 2019 (COVID-19) and responsible for the current pandemic. Recent SARS-CoV-2 susceptibility and transmission studies in cats show that the virus can replicate in these companion animals and transmit to other cats. Here, we present an in-depth study of SARS-CoV-2 infection, associated disease and transmission dynamics in domestic cats. Six 4- to 5-month-old cats were challenged with SARS-CoV-2 via intranasal and oral routes simultaneously. One day post challenge (DPC), two sentinel contact cats were co-mingled with the principal infected animals. Animals were monitored for clinical signs, clinicopathological abnormalities and viral shedding throughout the 21 DPC observation period. *Postmortem* examinations were performed at 4, 7 and 21 DPC to investigate disease progression. Viral RNA was not detected in blood but transiently in nasal, oropharyngeal and rectal swabs and bronchoalveolar lavage fluid as well as various tissues. Tracheobronchoadenitis of submucosal glands with the presence of viral RNA and antigen was observed in airways of the infected cats on 4 and 7 DPC. Serology showed that both, principal and sentinel cats, developed SARS-CoV-2-specific and neutralizing antibodies to SARS-CoV-2 detectable at 7 DPC or 10 DPC, respectively. All animals were clinically asymptomatic during the course of the study and capable of transmitting SARS-CoV-2 to sentinels within 2 days of comingling. The results of this study are critical for our understanding of the clinical course of SARS-CoV-2 in a naturally susceptible host species, and for risk assessment of the maintenance of SARS-CoV-2 in felines and transmission to other animals and humans.

## 1. Introduction

Coronaviruses are enveloped single-stranded, positive-sense RNA viruses that belong to the order *Nidovirales* in the family *Coronaviridae,* subfamily *Orthocoronavirinae,* and are comprised of four genera: *Alphacoronavirus, Betacoronavirus, Gammacoronavirus, and Deltacoronavirus* (Fehr and Perlman 2015). Many alpha- and betacoronaviruses originate from bats, while gamma- and deltacoronaviruses originate in birds (Woo, Lau et al. 2012). The Severe Acute Respiratory Syndrome-related Coronaviruses (SARS-CoV and SARS-CoV-2), and the Middle East Respiratory Syndrome coronavirus (MERS-CoV) belong to the genus betacoronavirus (Gorbalenya, Baker et al. 2020; Fung and Liu 2019). Alpha- and betacoronaviruses infect only mammals and cause important diseases of humans, cattle, pigs, cats, dogs, horses, and camels (Saif, 2004; Woo, Lau et al. 2012; Fehr and Perlman 2015). In general, coronaviruses cause respiratory, enteric, and systemic infections in humans and numerous animal hosts (Saif, 2004, Fehr and Perlman 2015). Importantly, coronaviruses can occasionally cross the species barriers (Drexler, Corman et al. 2014; Corman, Muth et al. 2018).

Bats have been identified as a reservoir species for many coronaviruses including those causing important human epidemics, namely SARS-CoV in 2002-2003 and MERS-CoV in 2012 (Drexler, Corman et al. 2014). Camels have since been shown to serve as the primary intermediate and reservoir host for MERS-CoV, causing continued zoonotic animal-to-human transmissions (de Wit, Doremalen et al. 2016). During the SARS-CoV epidemic, infected domestic cats were identified from households of SARS-CoV positive patients, and both cats and ferrets were subsequently experimentally shown to be easily infected and transmit SARS-CoV (Martina, Haagmans et al. 2003; van den Brand, Haagmans et al. 2008).

SARS-CoV-2 is the cause of Coronavirus Disease 2019 (COVID-19) and responsible for the current global pandemic (Zhou, Yang et al. 2020). A zoonotic transmission event amplified at a seafood and animal market in Wuhan, Hubei Province, China, is suspected to be the site of the first significant infectious outbreak in humans (Li, Guan et al. 2020), with bats and/or pangolins being speculated as the potential origin species based on the sequence homology of coronaviruses isolated from these animals (Anderson, Rambaut et al. 2020; Zhang, Wu et al. 2020; Zhou, Yang et al. 2020).

Since the outbreak of SARS-CoV-2 was first identified in December of 2019, it has been demonstrated that SARS-CoV-2 can naturally and experimentally infect several animal species (Lakdawala and Menachery 2020; Hernandez, Abad et al. 2020; Oreshkova et al. 2020). There have been multiple case reports of natural transmission of the virus from humans to dogs and cats, infection of “big cats” (i.e. a lion and tigers) at the Bronx Zoo, and mink on farms in The Netherlands, Denmark and Spain (Newman, Smith et al. 2020; Leroy, Gouilh et al. 2020; Oreshkova et al., 2020). In a recent animal susceptibility study, dogs, cats, ferrets, pigs, chickens and ducks were experimentally infected with SARS-CoV-2 (Shi, Wen et al. 2020). The results from that study show that both cats and ferrets were efficiently infected and could transmit the virus, dogs showed low susceptibility, while pigs and avian species were not permissive hosts. In addition, non-human primates (NHPs), hamsters and hACE2 transgenic or adenovirus transduced mice have also been evaluated as potential animal models for SARS-CoV-2 and seem to be susceptible showing mild to severe clinical signs (Cleary, Pitchford et al. 2020; Lakdawala and Menachery 2020).

The close association between humans and animals including companion animals, livestock and wildlife species, raises concerns regarding the potential risks of transmission of SARS-CoV-2 from humans to animals (“reverse zoonosis”), and the potential role infected animals could play in perpetuating the spread of the disease (Hernandez, Abad et al. 2020; Leroy, Gouilh et al. 2020). Further research of SARS-CoV-2 infection in various animal species is needed in order to identify susceptible hosts and to better understand the infection, disease, clinical course and transmission capabilities of susceptible animal species. This knowledge is important for risk assessment, implementing mitigation strategies, addressing animal welfare issues, and to develop preclinical animal models for evaluating drug and vaccine candidates for COVID-19.

Here we present an in-depth study of SARS-CoV-2 infection, associated disease and transmission in domestic cats. Clinical evaluation of weight, body temperature, blood parameters, serology, viral RNA shedding and RNA distribution in tissues and organ systems, and associated pathological findings are presented and discussed.

## 2. Material and methods

### 2.1. Cells and Virus

Vero E6 cells (ATCC^®^ CRL-1586™, American Type Culture Collection, Manassas, VA, USA) were used for virus propagation and titration. Cells were cultured in Dulbecco’s Modified Eagle’s Medium (DMEM, Corning, New York, N.Y, USA), supplemented with 5% fetal bovine serum (FBS, R&D Systems, Minneapolis, MN, USA) and antibiotics/antimycotics (Fisher Scientific, Waltham, MA, USA), and maintained at 37 °C under a 5% CO_2_ atmosphere. The SARS-CoV-2 USA-WA1/2020 strain was acquired from Biodefense and Emerging Infection Research Resources Repository (catalogue # NR-52281, BEI Resources, Manassas, VA, USA) and passaged 3 times in Vero E6 cells to establish a stock virus for inoculation of animals. This stock virus was sequenced by next generation sequencing (NGS) using the Illumina MiSeq and its consensus sequence was found to be 100% homologous to the original USA-WA1/2020 strain (GenBank accession: MN985325.1). To determine infectious virus titer, 10-fold serial dilutions were performed on 96-well plates of Vero E6 cells. The presence of cytopathic effect (CPE) after 96 hours incubation was used to calculate the 50% tissue culture infective dose (TCID_50_)/ml using the Spearman-Karber method (Hierholzer and Killington 1996). The prepared SARS-CoV-2 stock has a titer of 1 × 10^6^ TCID_50_/ml; this virus stock was used for experimental infection of the cats.

### 2.2. Animals and experimental design

#### 2.2.1. Ethics statement for use of animals

All animal studies and experiments were approved and performed under the Kansas State University (KSU) Institutional Biosafety Committee (IBC, Protocol #1460) and the Institutional Animal Care and Use Committee (IACUC, Protocol #4390) in compliance with the Animal Welfare Act. All animal and laboratory work were performed in biosafety level-3+ and -3Ag laboratory and facilities in the Biosecurity Research Institute at KSU in Manhattan, KS, USA.

#### 2.2.2. Virus challenge of animals

Ten 4.5- to 5-month old intact male cats were acclimated for seven days to BSL-3Ag biocontainment prior to experimental procedures with feed and water *ad libitum*. These were antibody profile defined/specific pathogen free (APD/SPF) animals with no detectable antibody titers to feline herpesvirus (rhinotracheitis), feline calicivirus, feline panleukopenia virus, feline coronaviruses, feline immunodeficiency virus, *Chlamydia felis* and *Toxoplasma gondii* obtained from Marshall BioResources (North Rose, New York, USA). The cats were placed into three groups (Figure 1, **and** Table 1). Group 1 (principal infected animals) consisted of six cats (three cats per housing unit), were inoculated simultaneously via the intranasal and oral routes with a total dose of 1 × 10^6^ TCID_50_ of SARS-CoV-2 in a total of 2 ml DMEM medium (0.5 ml per nostril and 1 ml oral). The cats in Group 2 (n=2; sentinel contact animals) and Group 3 (n=2; mock control animals) were housed in a separate room (Table 1, **and** Figure 1). Mock-infected cats (Group 3) were administered 2 ml DMEM via the intranasal and oral routes similar to Group 1 animals. At 1-day post challenge (DPC), the two cats in Group 2 were co-mingled with the principal infected animals in Group 1 (one cat per housing unit), and served as sentinel contact controls. The remaining two cats in Group 3 remained housed in a separate room and served as mock-infected negative controls. Principal infected animals were euthanized for *postmortem* examinations at 4 (n=2), 7 (n=2) and 21 (n=1) DPC to evaluate the course of disease. The two negative control animals in Group 3 were euthanized for *postmortem* examinations at 3 DPC. The remaining three animals from Group 1 (one principal infected animal) and Group 2 (two sentinel contact animals) were maintained for future re-infection studies, and not terminated as part of this study.

**Table 1.** Animal groups.

| **Table 1. Animal groups.** |  |  |  |
| --- | --- | --- | --- |
| Group | Treatment | Cat ID#s | Necropsy |
| 1 | Principal | 476, 493 | 4 DPC |
| 1 | Principal | 310, 597 | 7 DPC |
| 1 | Principal | 026 | 21 DPC |
| 1 | Principal | 328 | NA |
| 2 | Sentinel | 272, 903 | NA |
| 3 | Mock | 280, 403 | 3 DPC (not infected) |

NA=not available (enrolled in another study)

**Figure 1.**
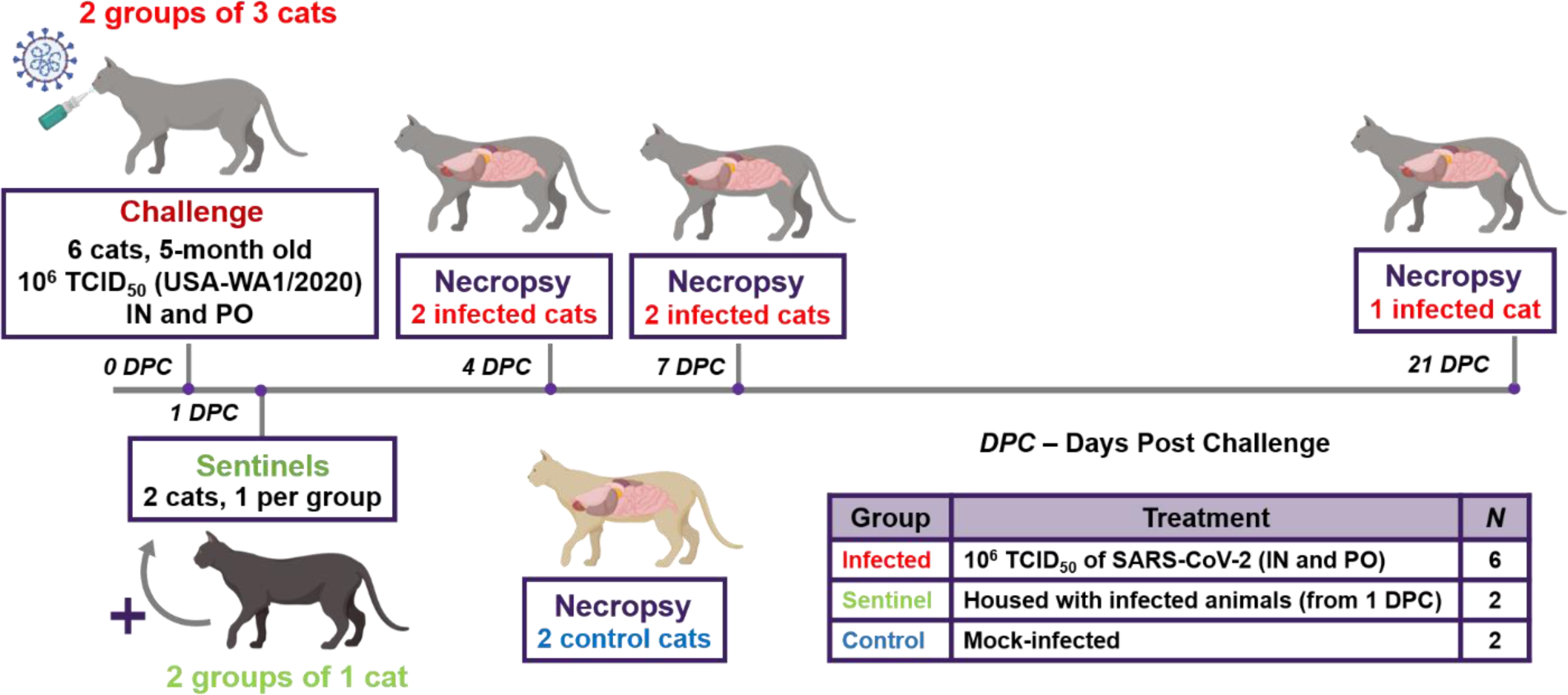
Study design. Ten cats were placed into three groups. Group 1 (principal infected animals) consisted of six cats (three cats/cage) and was inoculated via intranasal (IN) and oral (PO) routes simultaneously with a total dose of 1 × cephalic vein under anesthesia o 10^6^ TCID_50_ of SARS-CoV-2 in 2 ml DMEM. The cats in Group 2 (n=2; sentinel contact animals) and Group 3 (n=2; mock control animals) were housed in a separate room. At 1-day post challenge (DPC), the two cats in Group 2 were co-mingled with the principal infected animals in Group 1 (one cat per cage) and served as sentinel contact controls. The remaining two cats in Group 3 remained housed in a separate room and served as mock-infected negative controls. Principal infected animals were euthanized and necropsied at 4 (n=2), 7 (n=2) and 21 (n=1) DPC to evaluate the course of disease. The two negative control animals in Group 3 were euthanized and necropsied at 3 DPC. The remaining three animals from Group 1 (one principal infected animal) and Group 2 (two sentinel contact animals) were maintained for future re-infection studies.

#### 2.2.3. Clinical evaluations and sample collection

Cats were observed daily for clinical signs, such as: fever, anorexia, lethargy, respiratory distress, inappetence, depression, recumbency, coughing, sneezing, diarrhea/loose stool, vomiting and others. Weights of all cats were recorded on bleed days. Blood and serum were collected from all cats, including sentinel contact controls, on -1 DPC prior to infection, and on days 1, 3, 5, 7, 10, 14 and 21 DPC via venipuncture of the cephalic vein under anesthesia or during terminal bleeding by cardiac puncture. Nasal, oropharyngeal and rectal swabs were also collected on -1, 1, 3, 5, 7, 10, 14 and 21 DPC in 2 ml of virus transport medium (VTM; DMEM; Corning, New York, N.Y, USA) with antibiotics/antimycotic (Fisher Scientific, Waltham, MA, USA). Swabs were vortexed and supernatant aliquoted directly into cryovials or into RLT buffer (Qiagen, Germantown, MD, USA) and stored at -80 °C until further analysis.

A full *postmortem* examination was performed for each cat at the indicated time-points and gross changes (if any) were recorded. Tissues were collected either in 10% neutral-buffered formalin (Fisher Scientific, Waltham, MA, USA), or as fresh tissues which were then frozen at -80°C. A *postmortem* examination protocol was developed to collect the upper and lower respiratory tract, central nervous system (brain and cerebral spinal fluid [CSF) (**Figure S2**), gastrointestinal (GI) tract as well as accessory organs. The lungs were removed *in toto* including the trachea, and the main bronchi were collected at the level of the bifurcation and at the entry point into the lung lobe (**Figure S2B**). Lung lobes were evaluated based on gross pathology and collected and sampled separately (**Figure S2C**). Bronchoalveolar lavage fluid (BALF), nasal wash and urine were also collected during *postmortem* examination and stored at -80°C until analyzed. Fresh frozen tissue homogenates were prepared by thawing frozen tissue and placing 200 mg (± 50 mg) of minced tissue in a tube containing 1 ml DMEM culture medium and a steel bead (Qiagen, Germantown, MD, USA). Homogenization was performed with the TissueLyser LT (Qiagen, Germantown, MD, USA) for 30 seconds at 30 hertz repeated 3 times. Supernatant was retained after centrifugation for RNA extraction and quantitative reverse transcription real-time PCR (RT-qPCR).

### 2.3. Blood cell counts

Complete blood cell counts were performed using fresh EDTA blood samples run on an automated VetScan HM5 Hematology Analyzer (Abaxis, Inc., Union City, CA) according to the manufacturer’s recommended protocol using the VetScan HM5 reagent pack and recommended calibration controls. Blood cell analysis included: complete white blood cells, lymphocytes, monocytes, neutrophils, eosinophils, basophils, red blood cells, hematocrit, hemoglobin and platelets.

### 2.4. Serum biochemistry

Serum chemistry was performed using an automated VetScan VS2 Chemistry Analyzer (Abaxis, Inc., Union City, CA) according to the manufacturer’s recommended protocol. The Comprehensive Diagnostic Profile reagent rotor was used to perform complete chemistry and electrolyte analysis on 14 blood components: ALP, alkaline phosphatase; CRE, creatine; GLOB, globulin; PHOS, Phosphorous; GLU, glucose; BUN, blood urea nitrogen; Na, sodium; K, potassium; calcium; ALT, alanine aminotransferase; AMY, amylase; ALB, albumin; TBIL, total bilirubin; TP, total protein. Briefly, 100 µl of serum was added to the sample port of the reagent rotor, which was subsequently run in the machine.

### 2.5. RNA extraction and quantitative real-time reverse transcription PCR (RT-qPCR)

SARS-CoV-2-specific RNA was detected using a quantitative reverse transcription real time -PCR (RT-qPCR) assay. Briefly, tissue homogenates in VTM, blood, CSF, BALF, urine, and nasal, oropharyngeal and rectal swabs in VTM were mixed with an equal volume of RLT RNA stabilization/lysis buffer (Qiagen, Germantown, MD, USA), and 200µl of sample lysate was then used for extraction using a magnetic bead-based nucleic acid extraction kit (GeneReach USA, Lexington, MA) on an automated Taco™ mini nucleic acid extraction system (GeneReach USA, Lexington, MA) according to the manufacturer’s protocol with the following modifications: beads were added to the lysis buffer in the first well followed by the RLT sample lysate, then by the addition of 200 µl molecular grade isopropanol (ThermoFisher Scientific, Waltham, MA, USA), and finally, the last wash buffer B was replaced with 200 proof molecular grade ethanol (ThermoFisher Scientific, Waltham, MA, USA). Extraction positive controls (IDT, IA, USA; 2019-nCoV_N_Positive Control, diluted 1:100 in RLT) and negative controls were employed.

Quantification of SARS-CoV-2 RNA was performed using the N2 SARS-CoV-2 primer and probe sets (see: https://www.idtdna.com/pages/landing/coronavirus-research-reagents/cdc-assays) in a RT-qPCR protocol established by the CDC for the detection of SARS-CoV-2 nucleoprotein (N)-specific RNA (https://www.fda.gov/media/134922/download). This protocol has been validated in our lab for research use, using the qScript XLT One-Step RT-qPCR Tough Mix (Quanta BioSsciences, Beverly, MA, USA) on the CFX96 Real-Time thermocycler (BioRad, Hercules, CA, USA) using a 20-minute reverse transcription step and 45 cycle qPCR in a 20 µl reaction volume. A reference standard curve method using a 10-point standard curve of quantitated viral RNA (USA-WA1/2020 isolate) was used to quantify RNA copy number. RT-qPCR was performed in duplicate wells with a quantitated PCR positive control (IDT, IA, USA; 2019-nCoV_N_Positive Control, diluted 1:100) and four non-template control (NTC) on every plate. A positive Ct cut-off of 40 cycles was used. Data are presented as the mean and standard deviation of the calculated N gene copy number per ml of liquid sample or per mg of a 20% tissue homogenate.

### 2.6. Virus neutralizing antibodies

Virus neutralizing antibodies in sera were determined using microneutralization assay. Briefly, serum samples were initially diluted 1:10 and heat-inactivated at 56°C for 30 minutes while shaking. Subsequently, 100 µl per well of serum samples in duplicates were subjected to 2-fold serial dilutions starting at 1:20 through 1:2560 in 100 µl culture media. Then, 100 µl of 100 TCID_50_ of SARS-CoV-2 virus in DMEM culture media was added to 100 µl of the sera dilutions and incubated for 1 h at 37 °C. The 200 µl per well of virus sera mixture was then cultured on VeroE6 cells in 96-well plates. The corresponding SARS-CoV-2-negative cat sera, virus only and media only controls were also included in the assay. The neutralizing antibody titer was recorded as the highest serum dilution at which at least 50% of wells showed virus neutralization (NT_50_) based on the appearance of CPE observed under a microscope at 72 h post infection.

### 2.7. Detection of SARS-CoV-2 antibodies by indirect ELISA

To detect SARS-CoV-2 antibodies in sera, indirect ELISAs were performed with the recombinant viral proteins, nucleoprotein (N) and the receptor-binding domain (RBD), which were produced in-house. The N protein was produced in *E. coli* with a C-terminal His-Tag, and RBD was expressed in mammalian cells with a C-terminal Strep-Tag, and each were purified using either Ni-NTA (ThermoFisher Scientific, Waltham, MA, USA) or Strep-Tactin (IBA Lifesciences, Goettingen, Germany) columns, respectively, according to the manufacturers’ instructions. An optimal concentration of the respective coating antigen was first determined by checkerboard titration using SARS-CoV-2 positive and negative cat sera.

For indirect ELISAs, wells were coated with 100 ng of the respective protein in 100 µl per well coating buffer (Carbonate-bicarbonate buffer, catalogue number C3041, Sigma-Aldrich, St. Louis, MO, USA), then covered and incubated overnight at 4 °C. The next day, the plates were washed 2 times with phosphate buffered saline (PBS [pH=7.2-7.6]; catalogue number P4417, Sigma-Aldrich, St. Louis, MO, USA), blocked with 200 µl per well casein blocking buffer (Sigma-Aldrich, catalogue number B6429, St. Louis, MO, USA) and incubated for 1 h at room temperature. The plates were then washed 3 times with PBS-Tween20 (PBS-T; 0.5% Tween20 in PBS). Serum samples were pre-diluted 1:400 in casein blocking buffer, then 100 µl per well was added to the ELISA plate and incubated for 1 h at room temperature. Serial dilutions of the cat sera were not performed. The wells were washed 3 times with PBS-T, then 100 µl of Goat anti-Feline IgG (H+L) Secondary Antibody, HRP (ThermoFisher Scientific, catalogue number A18757, Waltham, MA, USA) diluted 1:2500 was added to each well and incubated for 1 h at room temperature. After 1 h, plates were washed 5 times with PBS-T, and 100 µl of TMB ELISA Substrate Solution (Abcam, catalogue number ab171525, Cambridge, MA, USA) was added to all wells of the plate. Following incubation at room temperature for 5 minutes, the reaction was stopped by adding 100 µl of 450 nm Stop Solution for TMB Substrate (Abcam, catalogue number ab171529, Cambridge, MA, USA) to all wells. The OD of the ELISA plates were read at 450 nm on an ELx808 BioTek plate reader (BioTek, Winooski, VT, USA). The cut-off for a sample being called positive was determined as follows: Average OD of negative serum + 3X standard deviation. Everything above this cut-off was considered as positive.

### 2.8. Gross pathology and histopathology

During *postmortem* examination, the head including the entire upper respiratory tract and central nervous system (brain), trachea and lower respiratory tract, lymphatic and cardiovascular systems GI and urogenital system, and integument were evaluated. CSF was collected with a syringe and needle via the atlanto-occipital (C0-C1) joint. Lungs were evaluated for gross pathology such as edema, congestion, discoloration, atelectasis, and consolidation. Tissue samples from the respiratory tract, nasal turbinates (rostral and deep), trachea (multiple levels; **Figure S2B**) and all 6 lung lobes (**Figure S2C**), GI (stomach, small and large intestine) and various other organs and tissues (spleen, kidney, liver, heart, tonsils, tracheo-bronchial and mesenteric lymph nodes, brain including olfactory bulb, and bone marrow) were collected and either fixed in 10% neutral-buffered formalin for histopathologic examination or frozen for RT-qPCR testing (see above section 2.2.3 and 2.5). Additional tissues placed in formalin for histological evaluation include the thymus, eye, salivary gland, skin (lip, eye lids, external nares, and ear), larynx, testes, adrenal gland, base of the tongue, urinary bladder and third eyelid. Tissues were fixed in formalin for 7 days then were transferred to 70% ethanol (ThermoFisher Scientific, Waltham, MA, USA) prior to trimming for embedding. Tissues were routinely processed and stained with hematoxylin and eosin following standard procedures within the histology laboratories of the Kansas State Veterinary Diagnostic Laboratory (KSVDL) and the Louisiana Animal Disease Diagnostic Laboratory (LADDL). Several veterinary pathologists independently examined slides and were blinded to the treatment groups.

### 2.9. SARS-CoV-2-specific RNAscope^®^ in situ hybridization (RNAscope^®^ ISH)

For RNAscope^®^ ISH, an anti-sense probe targeting the spike (S; nucleotide sequence: 21,563-25,384) of SARS-CoV-2, USA-WA1/2020 isolate (GenBank accession number MN985325.1) was designed (Advanced Cell Diagnostics [ACD], Newark, CA, USA) and used as previously described (Carossino et al. 2020). Four-micron sections of formalin-fixed paraffin-embedded tissues were mounted on positively charged Superfrost^®^ Plus Slides (VWR, Radnor, PA, USA). The RNAscope^®^ ISH assay was performed using the RNAscope 2.5 HD Red Detection Kit (ACD) as previously described (Carossino et al. 2019; Carossino et al. 2020). Briefly, deparaffinized sections were incubated with a ready-to-use hydrogen peroxide solution for 10 min at room temperature and subsequently subjected to Target Retrieval for 15 min at 98-102 °C in 1X Target Retrieval Solution. Tissue sections were dehydrated in 100% ethanol for 10 min and treated with Protease Plus for 20 min at 40 °C in a HybEZ™ oven (ACD). Slides were subsequently incubated with a ready-to-use probe mixture for 2 h at 40 °C in the HybEZ™ oven, and the signal amplified using a specific set of amplifiers (AMP1-6 as recommended by the manufacturer). The signal was detected using a Fast-Red solution (Red B: Red A in a 1:60 ratio) for 10 minutes at room temperature. Slides were counterstained with 50% Gill hematoxylin I (Sigma Aldrich, St Louis, MO, USA) for 2 min, and bluing performed with a 0.02% ammonium hydroxide in water. Slides were finally mounted with Ecomount^®^ (Biocare, Concord, CA, USA). Sections from mock- and SARS-CoV-2-infected Vero cell pellets were used as negative and positive assay controls.

### 2.10. SARS-CoV-2-specific immunohistochemistry (IHC)

For IHC, four-micron sections of formalin-fixed paraffin-embedded tissue were mounted on positively charged Superfrost^®^ Plus slides and subjected to IHC using a SARS-CoV-2-specific anti-nucleocapsid mouse monoclonal antibody (clone 6F10, BioVision, Inc., Milpitas, CA, USA) as previously described (Carossino et al., 2020). IHC was performed using the automated BOND-MAX and the Polymer Refine Red Detection kit (Leica Biosystems, Buffalo Grove, IL, USA), as previously described (Carossino et al. 2019). Following automated deparaffinization, heat-induced epitope retrieval (HIER) was performed using a ready-to-use citrate-based solution (pH 6.0; Leica Biosystems) at 100 °C for 20 min. Sections were then incubated with the primary antibody (diluted at 1 µg/ml in Antibody Diluent [Dako, Carpinteria, CA]) for 30 min at room temperature, followed by a polymer-labeled goat anti-mouse IgG coupled with alkaline phosphatase (30 minutes; Powervision, Leica Biosystems). Fast Red was used as the chromogen (15 minutes), and counterstaining was performed with hematoxylin. Slides were mounted with a permanent mounting medium (Micromount^®^, Leica Biosystems). Sections from mock- and SARS-CoV-2-infected Vero cell pellets were used as negative and positive assay controls.

## 3. Results

### SARS-CoV-2-infected domestic cats remain subclinical

Body temperature and clinical signs were recorded daily. No remarkable clinical signs were observed over the course of the study. At 2 DPC, some of the infected cats developed a temperature above 38.7 °C, but otherwise body temperature remained within the normal range for the remainder of the study. Body temperatures of sentinels were elevated at 1-, 10- and 12-days post contact (DPCo), but otherwise remained within normal range [**Figure S1A**]. Body weights of all cats increased throughout the study as expected for young animals without clinical disease [**Figure S1B**]. Complete blood counts and serum biochemistry were performed on days -1, 1, 3, 5, 7, 10, 14, and 21 DPC for the principal infected cats, and 2, 4, 6, 9, 13 and 20 DPCo for the sentinels. Overall, no significant changes in most blood cell parameters or serum biochemistry were observed. White blood cell (WBC) counts remained within normal limits for most animals during the course of the study; mildly increased WBC counts observed at -1 DPC and 1 DPC were attributed to stress early in the course of the study. No significant changes were observed in serum biochemical analytes except elevated alkaline phosphatase (ALP) levels in many animals starting at 5 DPC in the sentinels and after 7 DPC in the principal infected animals **[Figure S1C]**, which might indicate growth of subadult animals.

### SARS-CoV-2 RNA found throughout the respiratory tract

SARS-CoV-2 RNA was detected in nasal swabs of the principal infected cats at 1 through 10 DPC, with maximal quantities observed from 1 through 5 DPC (Figure 2A). The nasal swabs of contact animals became RNA positive for SARS-CoV-2 starting at day 2 DPCo (i.e. 3 DPC) and remained positive up to 9 DPCo/10 DPC, with a maximum on day 6 DPCo/7 DPC that is nearly as high as the copy number detected in the principal infected animals at 1 through 5 DPC (Figure 2A). The oropharyngeal swabs were RNA positive starting at 1 DPC through 10 DPC for the principals and 2 DPCo through 4 DPCo for the sentinels, with a maximum on 4 DPC and 4 DPCo, respectively (Figure 2B). Overall viral RNA in oropharyngeal swabs was approximately 1 to 2 logs lower than what was seen in the nasal swabs, for all cats except the principal infected at 4 DPC.

**Figure 2.**
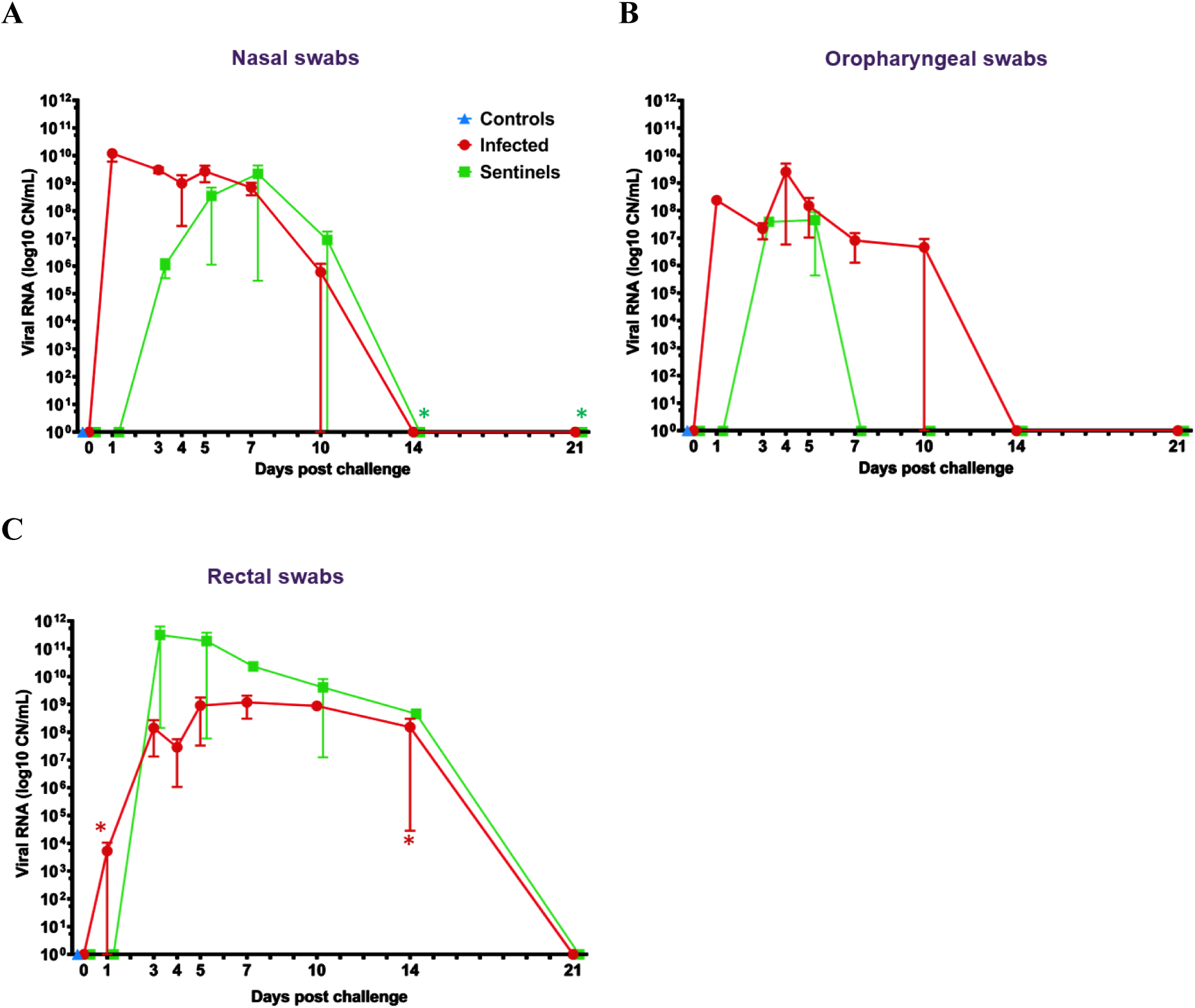
Felines shed SARS-CoV-2. RT-qPCR was performed on nasal (A), oropharyngeal (B) and rectal swabs (C) collected over the course of the 21-day study. Average viral copy number (CN) per mL with standard deviations are shown for each treatment group. Astrisks (*) indicate ½ of the sample RT-qPCR reactions were below the limit of detection.

Viral RNA was also detected in respiratory tract tissues in principal infected animals (Figure 3A, B). Fresh tissues collected during *postmortem* examination **[see Figure S2A, B, C]** from the nasal cavity, trachea, bronchi and all lung lobes were RNA positive for all animals at 4 and 7 DPC (Figure 3A, B), with RNA copy number ranging from 10^7^ to 10^11^ CN/mL for nearly all cats necropsied at these time points. Viral RNA levels in the lungs tended to be lower than the upper respiratory tract for cats necropsied at 7 DPC (Figure 3B). At 21 DPC viral RNA was detected within the upper respiratory samples, i.e. bronchi and right caudal lung lobe (Figure 3A, B). Nasal washes and BALF collected at necropsy from all principal infected cats examined at 4 and 7 DPC were RNA positive, but negative from the cat evaluated at 21 DPC (Table 2).

**Table 2.** Viral RNA (copy number/mL) detected in nasal washes, bronchoalveolar lung fluid (BALF), cerebrospinal fluid (CSF) and urine.

NA=not available (enrolled in another study)
|  |  |  |  |  |  |
| --- | --- | --- | --- | --- | --- |
| Cat ID# | DPC | Nasal washes | BALF | CSF | Urine |
| 476 | 4 | 6.5E+08 | 5.9E+08 | 2.5E+04* | 0 |
| 493 | 4 | 7.5E+08 | 3.2E+08 | 0 | 0 |
| 310 | 7 | 6.3E+07 | 2.0E+08 | 3.6E+06 | 0 |
| 597 | 7 | 5.6E+07 | 4.6E+05 | 0 | 0 |
| 026 | 21 | 0 | 0 | 0 | 0 |

\* 1/2 RT-qPCRs below limit of detection

**Figure 3.**
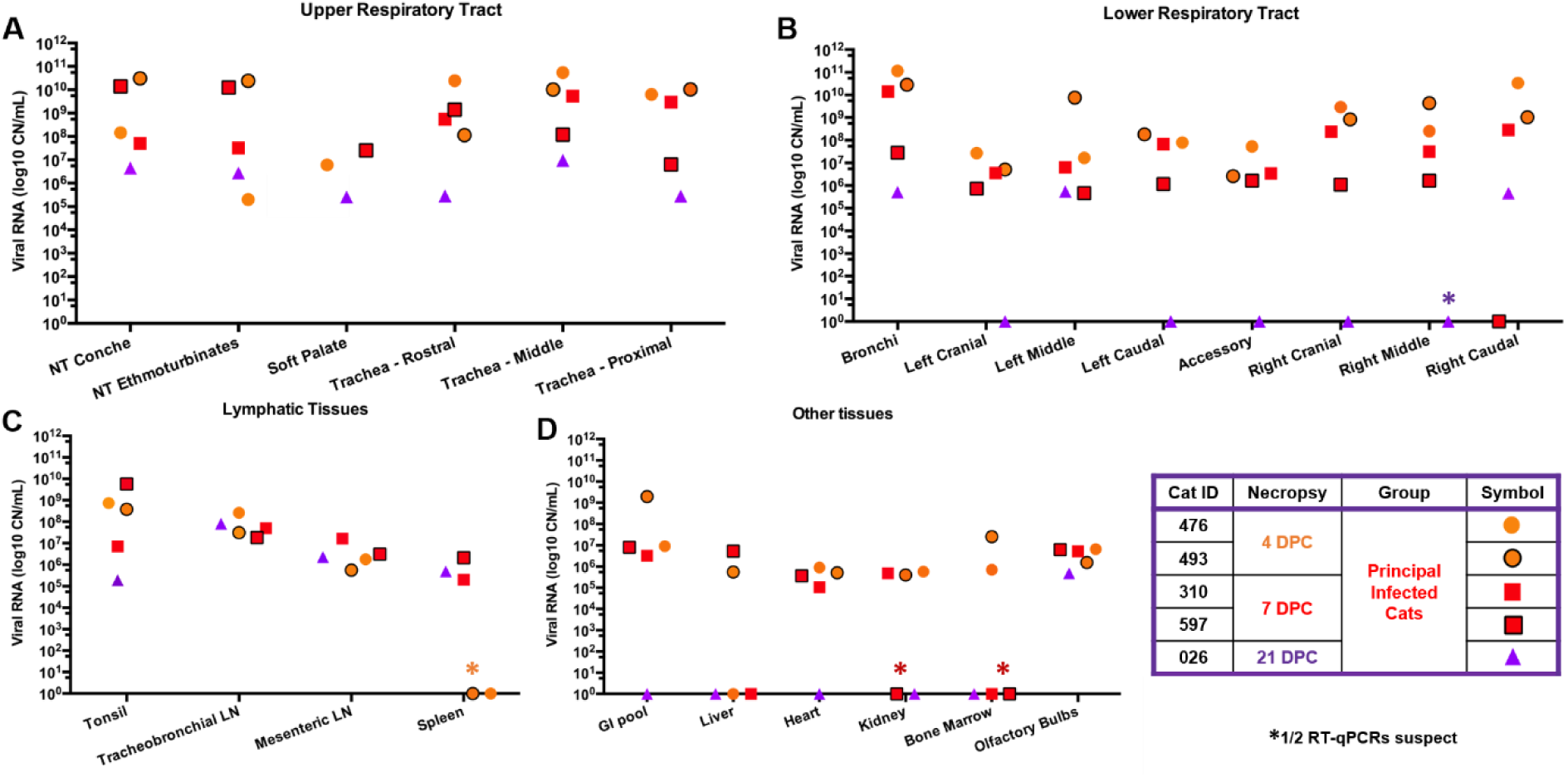
Viral RNA in tissues. SARS-CoV-2 copy number (CN) per mg tissues based on N is shown. LN=lymph node; GI=gastrointestinal

Gross pathology of the respiratory tract was accessed during *postmortem* examination. The lungs were removed *in toto* from each animal at 4, 7 and 21 DPC and demonstrated varying degree and distribution of edema, discoloration, congestion and atelectasis (**data not shown);** this could be attributed to euthanasia. Histologically, the pathological changes were limited to the upper and lower airways (larynx, trachea, and main, lobar and segmental bronchi of the lungs) of SARS-CoV-2 principal infected cats. Pathological findings are characterized by multifocal lymphocytic and neutrophilic tracheobronchoadenitis of seromucous lands of the lamina propria and submucosa of the trachea and bronchi. Changes range from minimal to mild at 4 DPC and progress to mild to moderate by 7 DPC (Figures 4, **and Figure S3**). Affected submucosal glands and associated ducts were variably distended (ectatic), lined by attenuated epithelium, contain various amounts of necrotic cell debris. In more severely affected foci glands are poorly delineated, and disrupted by mild to moderate numbers of infiltrating lymphocytes, macrophages and plasma cells, and few neutrophils (Figure 4, **and Figure S3**). No significant pathology was identified elsewhere in the pulmonary parenchyma (bronchioles, pulmonary vessels, alveolar spaces, alveolar septa and visceral pleura) of SARS-CoV-2-infected cats on 4 and 7 DPC. No significant histologic changes were noted in the respiratory tract at 21 DPC, with the submucosal architecture of the trachea and bronchi being unremarkable and within normal limits (Figure 4, **and Figure S3**).

**Figure 4.**
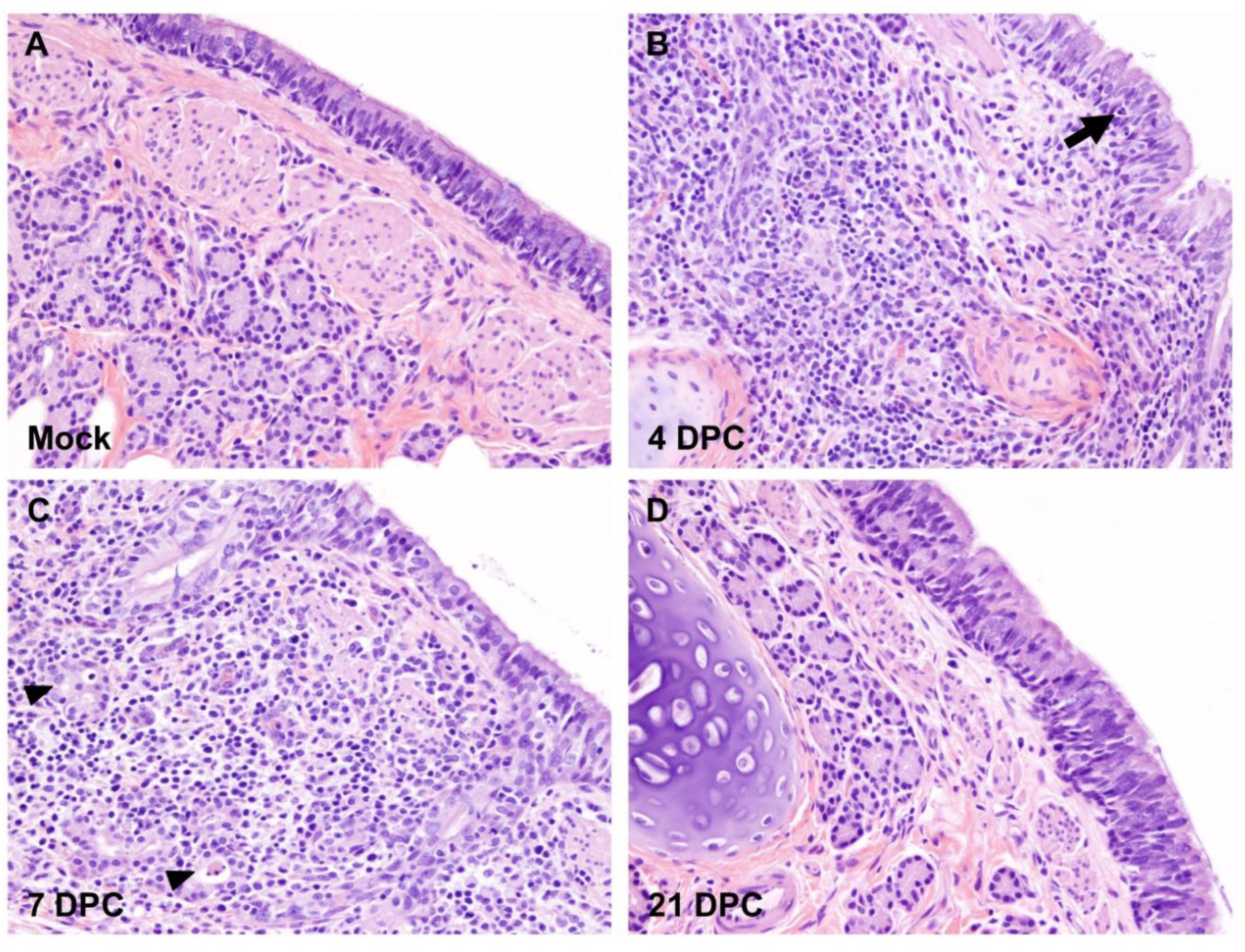
Histopathology of bronchi. Histological findings in the main bronchi of mock (A) and SARS-CoV-2 experimentally infected (B-D) cats. Histologic changes and their progression are similar to those observed in the trachea, with multifocal, widespread, mild to moderate lymphocytic and neutrophilic adenitis noted at 4 days post-challenge (DPC; B) and 7 DPC (C). Necrotic debris within distorted submucosal glands are indicated with arrowheads (C), and few transmigrating lymphocytes are indicated with an arrow (B). No histologic changes are noted at 21 DPC (D). H&E. Total magnification: 200X

The cellular tropism, distribution and abundance of SARS-CoV-2 were also investigated via the detection of viral RNA and viral antigen by RNAscope^®^ ISH and IHC, respectively. Presence of viral RNA and antigen correlated with the histological changes observed in the airways and were detected within epithelial cells of submucosal glands and associated ducts at 4 and 7 DPC (Figure 5, **and Figure S4**). SARS-CoV-2-positive submucosal glands were more frequently observed at 4 DPC compared to 7 DPC (Figure 5, **and Figure S4**). Viral RNA and viral antigen were not detected at 21 DPC (Figure 5, and **Figure S4**). Noteworthy, no viral RNA or antigen were detected within lining epithelial cells or elsewhere in the pulmonary parenchyma, including smaller airways (i.e. bronchioles) and alveoli, at 4, 7 or 21 DPC.

**Figure 5.**
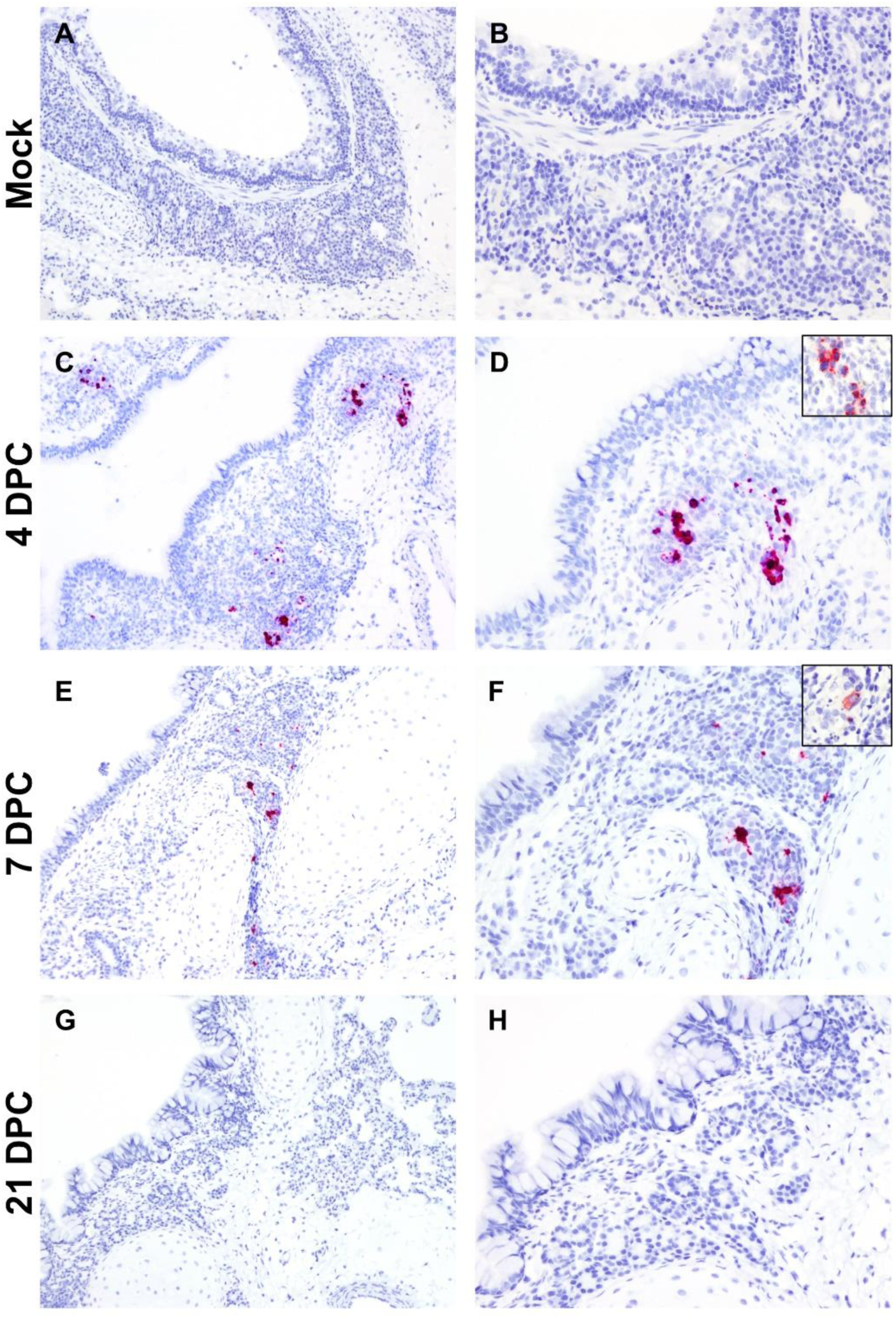
SARS-CoV-2 RNA and antigen detection in bronchi. SARS-CoV-2 tropism in bronchi of mock (A and B) experimentally (C-H) infected cats determined by S-specific RNAscope^®^ *in situ* hybridization (Fast Red) and anti-N-specific immunohistochemistry (IHC; Fast Red). The viral tropism is limited to glandular and ductular epithelial cells of multifocal, scattered submucosal glands, as noted in the trachea. Viral RNA is detected within infected cells at 4 days post-challenge (DPC; C and D) and, to a lower degree at 7 DPC (E and F). Few scattered glandular epithelial cells are positive for SARS-CoV-2 N antigen by IHC (D and F, insets). No viral RNA or antigen is detected at 21 DPC (G and H). Total magnification: 100X (A, C, E and G) and 200X (B, D, F, H).

### SARS-CoV-2 RNA found throughout non-respiratory organs and tissues

SARS-CoV-2 RNA was detected in rectal swabs starting at 3 DPC for principal infected cats and 2 DPCo for sentinel cats and were found positive up to 14 DPC or 13 DPCo, respectively. High level of RNA shedding was maintained from 3 DPC or 2 DPCo throughout 14 DPC or 13 DPCo before animals became RNA negative by 21 DPC or 20 DPCo for both principal infected and sentinel cats (Figure 2C). Urine collected directly from the bladder during *postmortem* examination of cats sacrificed at 4, 7, 21 DPC was negative by RT-qPCR (Table 2). Viral RNA was also detected in the GI tract and other organs and tissues in principal infected animals, such as tonsils, spleen, lymph nodes, kidney, liver, heart, bone marrow and olfactory bulb (Figure 3C, D). Tonsils, lymph nodes and olfactory bulbs of all cats were positive on 4, 7, and 21 DPC. The highest CN of RNA was detected in the tonsils, tracheobronchial and mesenteric lymph nodes at all days tested. Spleen was negative on 4 DPC, but all animals were positive on 7 and 21 DPC. RNA was present in the pooled tissue from the GI tract and heart in all animals on 4 and 7 DPC. Liver, kidney, and bone marrow were occasionally positive on different time points post infection. CSF was RT-qPCR positive from 1 of the 2 cats necropsied at 4 DPC and 1 of 2 cats necropsied at 7 DPC, but not at 21 DPC (Figure 3D, and Table 2). Blood from all cats collected at -1, 1, 3, 5, 7, 10, 14 and 21 DPC were RT-qPCR negative for SARS-CoV-2. Gross evaluation of the GI tract, the cardiovascular and central nervous system as well as additional visceral organs and lymphoid tissues revealed no macroscopic changes between SARS-CoV-2 and mock-infected cats at any time-point post infection. Additionally, no histopathological changes were identified in multiple segments of the GI tract or in other organs examined so far.

### Seroconversion of cats after SARS-CoV-2 infection

Sera collected at -1, 3, 5, 7, 10, 14 and 21 DPC was tested for the presence of SARS-CoV-2 specific antibodies. Virus neutralizing antibodies were detected in sera from all principal infected and contact sentinel cats at 10, 14 and 21 DPC, with neutralizing titers ranging from 1:40 to 1:320 (Table 3). Sera from principal or sentinel animals tested before 7 or 10 DPC were negative for neutralizing antibodies, respectively. Antibodies against the N protein were detected in all principal-infected cats starting at 7 DPC and in the two sentinel cats after 9 DPCo throughout the end of the observation period of 21 DPC (Table 3). Similarly, antibodies against the receptor binding domain protein were detected in all principal infected starting at 7 DPC and in the sentinel cats starting at 13 DPCo throughout the end of the observation period of 21 DPC.

**Table 3.** Felines develop SARS-CoV-2 specific and virus neutralizing antibodies

|  |  |  |  |  |  |  |  |  |  |  |  |  |  |  |  |  |
| --- | --- | --- | --- | --- | --- | --- | --- | --- | --- | --- | --- | --- | --- | --- | --- | --- |
| Anti-N antibodies |  |  |  |  |  | Anti-RBD antibodies |  |  |  |  | Virus neutralizing antibodies (NT<sub>50</sub>) |  |  |  |  |  |
| DPC | 5 | 7 | 10 | 14 | 21 | 5 | 7 | 10 | 14 | 21 | 1 | 5 | 7 | 10 | 14 | 21 |
| *Principals* |  |  |  |  |  |  |  |  |  |  |  |  |  |  |  |  |
| 026 | + | + | + | + | + | - | + | + | + | + | 0 | 0 | 1:80 | 1:320 | 1:80 | 1:40 |
| 328 | - | + | + | + | NA | + | + | + | + | NA | 0 | 0 | 1:20 | 1:160 | 1:160 | 1:160 |
| *Sentinels* |  |  |  |  |  |  |  |  |  |  |  |  |  |  |  |  |
| 272 | - | - | - | + | + | - | - | - | - | + | 0 | 0 | 0 | 1:160 | 1:160 | 1:160 |
| 903 | - | +/- | +/- | + | + | - | - | - | + | + | 0 | 0 | 0 | 1:80 | 1:160 | 1:160 |

NA=not available; N=nucleoprotein; RBD=receptor binding domain of spike protein; +/-=OD close to cut-off

## 4. Discussion

Identifying susceptible species and their capacity to transmit SARS-CoV-2 is critical for the determination of likely sources of reverse zoonosis and for mitigating viral spread. The development of animal models for COVID-19 is equally critical for studying the mechanisms of the disease and for evaluating the efficacy of potential vaccines, antiviral drugs and therapies. In this study we explored in-depth the infection, associated disease and transmission dynamics in 4- to 5-months old domestic cats. The cats were antibody profile defined/specific pathogen free (APD/SPF) animals with no antibody titers against various feline virus infections including feline coronaviruses. The detection of high levels of viral RNA from swab samples and in various organs and tissues, along with mild to moderate histologic changes in trachea and bronchi associated with viral RNA and viral antigen, and the development of SARS-CoV-2-specific antibodies demonstrates that cats were productively infected, without developing any obvious clinical signs. Viral RNA was detected at 4 and 7 DPC throughout the upper and lower respiratory tracts, GI tract, olfactory bulb, and lymphoid and other tissues of all inoculated cats; viral RNA was mostly cleared by 21 DPC in the lungs, BALF, nasal washes and other tissues including GI tract, but persisted in the upper respiratory tract, and lymphoid tissue. In contrast, viral RNA was detected in the spleen only at 7 and 21 DPC, but not at 4 DPC. Absence of viral RNA shedding and histologic changes in the lungs and trachea at 21 DPC as well as seroconversion suggests that the cats were responding to experimental SARS-CoV-2 infection by mounting a humoral immune response and recovering from the infection at 3-weeks post experimental inoculation. Furthermore, the principal infected cats were able to transmit the virus to negative contact animals within 2 days of contact housing similar as reported by Halfmann and coworkers (2020). High amounts of viral RNA shedding through the respiratory and GI tract are most likely responsible for the transmission to the sentinel animals. Shi and coworkers (2020) determined that airborne transmission of SARS-CoV-2 among cats is possible but not highly effective.

Our results, the studies by Shi et al. (2020) and Halfmann et al. (2020), as well as the reports of pet cats in households with COVID-19 patients (Newman, Smith et al. 2020; Leroy, Gouilh et al. 2020), show that felines are susceptible to SARS-CoV-2 infection and could be potential virus reservoirs. Consistent with our results, Shi et al. (2020) and Halfmann et al. (2020) also reported no obvious clinical signs in SARS-CoV-2 infected cats which were older than 4 months. We detected high viral RNA levels throughout nearly all tissues tested for all cats at 4 and 7 DPC, with reduced levels or clearing by 21 DPC in some tissues: residual RNA was detected mainly in the upper respiratory tract, the lymphoid tissues and the CNS. Since no virus was detected in blood, it remains to be studied how the virus reaches and infects non-respiratory tissues. Shi and colleagues (2020) also reported that viral RNA and infectious virus was detected throughout upper and lower respiratory tracts in juvenile (70-100 days old) and subadult cats (6-9 months old) at 3 DPC but it was cleared from most lung tissues of subadult cats by 6 DPC; however, in juvenile cats, viral RNA and infectious virus was still present on 6 DPC in the lower respiratory tract (Shi et al., 2020). In contrast, no viral RNA or virus was detected in other organs of any of these cats which included brain, heart, submaxillary lymph nodes, kidney, spleen, liver, and pancreas at 3 or 6 DPC (Shi, Wen et al. 2020). Similar to Shi and colleagues (2020) who detected viral RNA and infectious virus in the small intestine of most of the animals we found shedding of viral RNA in rectal swabs up to 14 DPC and viral RNA in pooled GI tract tissues on 4 and 7 DPC. In contrast, Halfmann and coworkers (2020) found all rectal swabs to be virus negative. These differences may be explained by the age of the cats used in these studies, and the different virus strains used. Shi et al. (2020) were also describing SARS-CoV-2 associated pathological changes in juvenile and subadult cats, and noticed more severe pathology was associated with the juvenile cats, including histopathological lesions in nasal and tracheal mucosal epithelia and lungs (Shi, Wen et al. 2020). Our results are consistent with that study, showing cats 4 to 5 months of age had mild to moderate histologic alterations identified as tracheobronchoadenitis within the airways. The macroscopic lung lesions observed at *postmortem* examinations were most likely due to the euthanasia with barbiturates. Importantly, all SARS-CoV-2 infected cats (principal and sentinel animals) in our study mounted an antiviral and neutralizing antibody response during the 21-day observation period. Virus specific antibodies to the N and RBD proteins were detected in all principal animals starting at 7 DPC as well as virus neutralizing antibodies. Other studies reported detection of IgG antibodies against RBD as early as 1 DPC (Halfmann, Hatta et al. 2020), and virus specific and neutralizing antibodies in all principal inoculated cats, but only in 2 out of the 6 sentinel animals (Shi, Wen et al. 2020). None of these studies detected virus or viral RNA in the blood.

Similar to previous experimental infection studies with SARS-CoV-1 (Martina, Haagmans et al. 2003; van den Brand et al. 2008) and more recently SARS-CoV-2 (Shi, Wen et al. 2020) in cats, histological changes within the airways (namely trachea and main, lobar and segmental bronchi) under our experimental infection conditions with SARS-CoV-2 in cats are limited to mild to moderate neutrophilic and lymphocytic tracheobronchoadenitis with associated intralesional detection of viral RNA and viral antigen. While SARS-CoV-1 antigen was identified in sporadic tracheal and bronchial epithelial cells in a single experimentally infected cat (Martina, Haagmans et al. 2003; van den Brand, Haagmans et al. 2008), we determined that the airway epithelial lining of the trachea and the main, lobar and segmental bronchi seems non-permissive to SARS-CoV-2 replication *in vivo* at least in subadult cats (4- to 5-months old) as demonstrated by the lack of viral RNA and viral antigen within these specific cells using RNAscope^®^ ISH and IHC; this correlated with the lack of histologic alterations on the surface epithelium other than sporadic neutrophil transmigration. These findings are in partial agreement to those of another recent study on the susceptibility of cats to SARS-CoV-2 infection (Shi, Wen et al. 2020), where mild histologic alterations in the tracheal lumen and epithelium were reported in the absence of detectable viral antigen. Interestingly, and in contrast with reports of SARS-CoV-1 infection in cats, SARS-CoV-2 infected 4- to 5-months old cats did not demonstrate histological changes within the small airways or the pulmonary parenchyma consistent with interstitial pneumonia or diffuse alveolar damage (DAD)/acute respiratory distress syndrome (ARDS), such as inflammatory infiltrates within alveolar septa or alveolar spaces, intra-alveolar fibrin or hyaline membranes, or pneumocyte type II hyperplasia. In addition, there is no evidence of SARS-CoV-2 infection within pneumocytes or alveolar macrophages as demonstrated by the absence of viral RNA (RNAscope^®^ ISH) and viral antigen (IHC). These findings correlate with the absence of clinically evident respiratory disease following experimental infection, with the duration and magnitude of viral shedding, and with the onset of SARS-CoV-2 specific antibody responses; and with no histologic changes or viral RNA and viral antigen present within the respiratory tissues by 21 DPC. While a recent study evaluating the susceptibility of domestic cats to SARS-CoV-2 (Shi et al., 2020) suggested the occurrence of additional histologic alterations in the pulmonary vasculature and alveolar spaces, none of these changes were noted in our experimental model. Additional studies are needed to determine whether these differences are due to the breed and age of the domestic cats, the virus isolate used for infection or other factors.

Together these findings warrant COVID-19 screening of felines for surveillance/epidemiological purposes and for implementing of mitigation strategies; they also point towards nasal swabs/washes and rectal swabs as appropriate diagnostic samples. This information will be important for providing appropriate veterinary care for infected cats and other cats in their surroundings, for protection of veterinary personnel, animal caretakers and pet owners, and for implementing quarantine measures to prevent transmission between felines, people and potentially other susceptible animals. The ease of transmission between domestic cats indicates a significant public health necessity to investigate the potential chain of human-cat-human transmission potential. It is also critical that pet owners are educated on the risks and preventative measures in order to calm fears and discourage animal abandonment.

Although asymptomatic, cats can be productively infected and readily transmit SARS-CoV-2 to other susceptible cats, and thus may serve as potential models for asymptomatic COVID-19 infections in humans. It could also offer a viable model for testing vaccines and antiviral candidates for companion animals and for drugs with a problematic pharmacokinetic profile in rodents, ferrets or nonhuman primates. However, a preclinical animal model that represents the clinical symptoms and disease observed in severe COVID-19 patients is still needed to improve evaluation of vaccines, antiviral drugs and other therapies.

Further research is needed to adapt models to recapitulate severe disease observed in humans. One area to explore is the effect of age on associated disease and recovery. Only cats less than 1 year old were evaluated in this study and the studies by Shi et al. (2020) and Halfmann et al. (2020), but what SARS-CoV-2 infection looks like in adult and older aged cats, as well as, if re¬infection of cats can occur and what re-infection looks like was not explored in these studies. Recent non peer-reviewed work by Bosco-Lauth and colleagues (2020) investigated experimental SARS-CoV-2 challenge and direct transmission in adult cats 5 to 8 years old, and re-infection after 28 DPC. The results from that study demonstrate that adult cats become infected without clinical signs with pathology limited to respiratory airways, and can readily transmit the virus to naïve cats. Furthermore, the adult cats appear to be protected from reinfection and mounted significantly higher neutralizing antibody responses compared to findings from our study and by Shi et al. (2020). Studies to better understand the mechanisms of infection and the range of symptoms and pathology associated with SARS-CoV-2 in various preclinical models of COVID-19 are critical for the development of vaccines and treatments of this disease.

## Acknowledgments

We gratefully thank the staff of KSU Biosecurity Research Institute, the histological laboratory at the Kansas State Veterinary Diagnostic Laboratory (KSVDL), members of the Histology and Immunohistochemistry sections at the Louisiana Animal Disease Diagnostic Laboratory (LADDL), the CMG staff and Gleyder Roman-Sosa, Yonghai Li, Emily Gilbert-Esparza, Chester McDowell at KSU, and Drs. James MacLachlan and Dennis Wilson at the School of Veterinary Medicine, University of California, Davis for their expert pathology consultations. The following reagent was obtained through BEI Resources, National Institute of Allergy and Infectious Diseases (NIAID), National Institutes of Health (NIH): SARS-CoV-2 Virus strain USA-WA1/2020 (catalogue # NR-52281).

## Funding

Funding for this study was provided through grants from NBAF Transition Funds, the NIAID Centers of Excellence for Influenza Research and Surveillance under contract number HHSN 272201400006C and the Department of Homeland Security Center of Excellence for Emerging and Zoonotic Animal Diseases under grant no. 2010-ST061-AG0001 to JAR. This study was also partially supported by the Louisiana State University, School of Veterinary Medicine start-up fund (PG 002165) to UBRB and the U.S. Department of Agriculture, Agricultural Research Service (58-32000-009-00D) to WCW, by the Center for Research for Influenza Pathogenesis (CRIP), a NIAID supported Center of Excellence for Influenza Research and Surveillance (CEIRS, contract # HHSN272201400008C), and by the generous support of the JPB Foundation, the Open Philanthropy Project (research grant 2020-215611 (5384)) and anonymous donors to AG-S.

Mention of trade names or commercial products in this publication is solely for the purpose of providing specific information and does not imply recommendation or endorsement by the U.S. Department of Agriculture. USDA is an equal opportunity provider and employer.

## References

Andersen, K. G., A. Rambaut, W. I. Lipkin, E. C. Holmes and R. F. Garry (2020). “The proximal origin of SARS-CoV-2.” Nat Med 26(4): 450–452.

Bosco-Lauth, A. M., A. E. Hartwig, S. M. Porter, P. W. Gordy, M. Nehring, A. D. Byas, S. VandeWoude, I. K. Ragan, R. M. Maison, R. A. Bowen. “Pathogenesis, transmission and response to re-exposure of SARS-CoV-2 in domestic cats.” bioRxiv 2020.05.28.120998; doi: https://doi.org/10.1101/2020.05.28.120998

Carossino, M., P. Dini, T. S. Kalbfleisch, A. T. Loynachan, I. F. Canisso, R. F. Cook, P. J. Timoney and U. B. R. Balasuriya (2019). “Equine arteritis virus long-term persistence is orchestrated by CD8+ T lymphocyte transcription factors, inhibitory receptors, and the CXCL16/CXCR6 axis.” PLoS Pathos 15(7): e1007950.

Carossino M., Ip H. S., Richt J. A., Shultz K., Harper K., Loynachan A. T., Del Piero F., Balasuriya U. B. R. (2020) “Detection of SARS-CoV-2 by RNAscope^®^ in situ Hybridization and Immunohistochemistry Techniques.” Arch Virol *In Press* DOI: 10.1007/s00705-020-04737-w

Cleary, S. J., S. C. Pitchford, R. T. Amison, R. Carrington, C. L. Robaina Cabrera, M. Magnen, M. R. Looney, E. Gray and C. P. Page (2020). “Animal models of mechanisms of SARS-CoV-2 infection and COVID-19 pathology.” Br J Pharmacol 10.1111/bph. 15143. Advance online publication. https://doi.org/10.1111/bph.15143

Corman, V. M., D. Muth, D. Niemeyer and C. Drosten (2018). “Hosts and Sources of Endemic Human Coronaviruses.” Adv Virus Res 100: 163–188.

de Wit, E., N. van Doremalen, D. Falzarano and V. J. Munster (2016). “SARS and MERS: recent insights into emerging coronaviruses.” Nature Reviews Microbiology 14(8): 523–534.

Drexler, J. F., V. M. Corman and C. Drosten (2014). “Ecology, evolution and classification of bat coronaviruses in the aftermath of SARS.” Antiviral Res 101: 45–56.

Fehr, A. R. and S. Perlman (2015). “Coronaviruses: an overview of their replication and pathogenesis.” Methods Mol Biol 1282: 1–23.

Fung, T. S. and D. X. Liu (2019). “Human Coronavirus: Host-Pathogen Interaction.” Annu Rev Microbiol 73: 529–557.

Gorbalenya, A.E., Baker, S.C., Baric, R.S. et al. (2020) “The species Severe acute respiratory syndrome-related coronavirus: classifying 2019-nCoV and naming it SARS-CoV-2.” Nat Microbiol 5: 536–544.https://doi.org/10.1038/s41564-020-0695-z

Halfmann, P. J., M. Hatta, S. Chiba, T. Maemura, S. Fan, M. Takeda, N. Kinoshita, S. I. Hattori, Y. Sakai-Tagawa, K. Iwatsuki-Horimoto, M. Imai and Y. Kawaoka (2020). “Transmission of SARS-CoV-2 in Domestic Cats.” N Engl J Med.

Hernandez, M., D. Abad, J. M. Eiros and D. Rodriguez-Lazaro (2020). “Are Animals a Neglected Transmission Route of SARS-CoV-2?” Pathogens 9(6).

Hierholzer, J. C. and R. A. Killington (1996). 2 - Virus isolation and quantitation A2 - Mahy, Brian WJ. Virology Methods Manual. H. O. Kangro. London, Academic Press: 25–46.

Lakdawala, S. S. and V. D. Menachery (2020). “The search for a COVID-19 animal model.” Science 368(6494): 942–943.

Leroy, E. M., M. Ar Gouilh and J. Brugere-Picoux (2020). “The risk of SARS-CoV-2 transmission to pets and other wild and domestic animals strongly mandates a one-health strategy to control the COVID-19 pandemic.” One Health: 100133.

Li, Q., X. Guan, P. Wu, X. Wang, L. Zhou, Y. Tong, R. Ren, K. S. M. Leung, E. H. Y. Lau, J. Y. Wong, X. Xing, N. Xiang, Y. Wu, C. Li, Q. Chen, D. Li, T. Liu, J. Zhao, M. Liu, W. Tu, C. Chen, L. Jin, R. Yang, Q. Wang, S. Zhou, R. Wang, H. Liu, Y. Luo, Y. Liu, G. Shao, H. Li, Z. Tao, Y. Yang, Z. Deng, B. Liu, Z. Ma, Y. Zhang, G. Shi, T. T. Y. Lam, J. T. Wu, G. F. Gao, B. J. Cowling, B. Yang, G. M. Leung and Z. Feng (2020). “Early Transmission Dynamics in Wuhan, China, of Novel Coronavirus-Infected Pneumonia.” N Engl J Med 382(13): 1199–1207.

Martina, B. E., B. L. Haagmans, T. Kuiken, R. A. Fouchier, G. F. Rimmelzwaan, G. Van Amerongen, J. S. Peiris, W. Lim and A. D. Osterhaus (2003). “Virology: SARS virus infection of cats and ferrets.” Nature 425(6961): 915.

Newman, A., D. Smith, R. R. Ghai, R. M. Wallace, M. K. Torchetti, C. Loiacono, L. S. Murrell, A. Carpenter, S. Moroff, J. A. Rooney and C. Barton Behravesh (2020). “First Reported Cases of SARS-CoV-2 Infection in Companion Animals - New York, March-April 2020.” MMWR Morb Mortal Wkly Rep 69(23): 710–713.

Oreshkova, N., R. J. Molenaar, S. Vreman, F. Harders, B. B. Oude Munnink, R. W. Hakze-van der Honing, N. Gerhards, P. Tolsma, R. Bouwstra, R. S. Sikkema, M. G. Tacken, M. M. de Rooij, E. Weesendorp, M. Y. Engelsma, C. J. Bruschke, L. A. Smit, M. Koopmans, W. H. van der Poel and A. Stegeman (2020). “SARS-CoV-2 infection in farmed minks, the Netherlands, April and May 2020.” Euro Surveill 25(23).

Saif, L. J. (2004). “Animal coronaviruses: what can they teach us about the severe acute respiratory syndrome?” Rev Sci Tech 23(2): 643–660.

Shi, J., Z. Wen, G. Zhong, H. Yang, C. Wang, B. Huang, R. Liu, X. He, L. Shuai, Z. Sun, Y. Zhao, P. Liu, L. Liang, P. Cui, J. Wang, X. Zhang, Y. Guan, W. Tan, G. Wu, H. Chen and Z. Bu (2020). “Susceptibility of ferrets, cats, dogs, and other domesticated animals to SARS-coronavirus 2.” Science 368(6494): 1016–1020.

van den Brand, J. M., B. L. Haagmans, L. Leijten, D. van Riel, B. E. Martina, A. D. Osterhaus and T. Kuiken (2008). “Pathology of experimental SARS coronavirus infection in cats and ferrets.” Vet Pathol 45(4): 551–562.

Woo, P. C., S. K. Lau, C. S. Lam, C. C. Lau, A. K. Tsang, J. H. Lau, R. Bai, J. L. Teng, C. C. Tsang, M. Wang, B. J. Zheng, K. H. Chan and K. Y. Yuen (2012). “Discovery of seven novel Mammalian and avian coronaviruses in the genus deltacoronavirus supports bat coronaviruses as the gene source of alphacoronavirus and betacoronavirus and avian coronaviruses as the gene source of gammacoronavirus and deltacoronavirus.” J Virol 86(7): 3995–4008.

Zhang, T., Q. Wu and Z. Zhang (2020). “Pangolin homology associated with 2019-nCoV.” bioRxiv: 2020.2002.2019.950253.

Zhou, P., X. L. Yang, X. G. Wang, B. Hu, L. Zhang, W. Zhang, H. R. Si, Y. Zhu, B. Li, C. L. Huang, H. D. Chen, J. Chen, Y. Luo, H. Guo, R. D. Jiang, M. Q. Liu, Y. Chen, X. R. Shen, X. Wang, X. S. Zheng, K. Zhao, Q. J. Chen, F. Deng, L. L. Liu, B. Yan, F. X. Zhan, Y. Y. Wang, G. F. Xiao and Z. L. Shi (2020). “A pneumonia outbreak associated with a new coronavirus of probable bat origin.” Nature 579(7798): 270–273.

