## Supplementary figures S1-4 for "SARS-CoV-2 infection, disease and transmission in domestic cats"

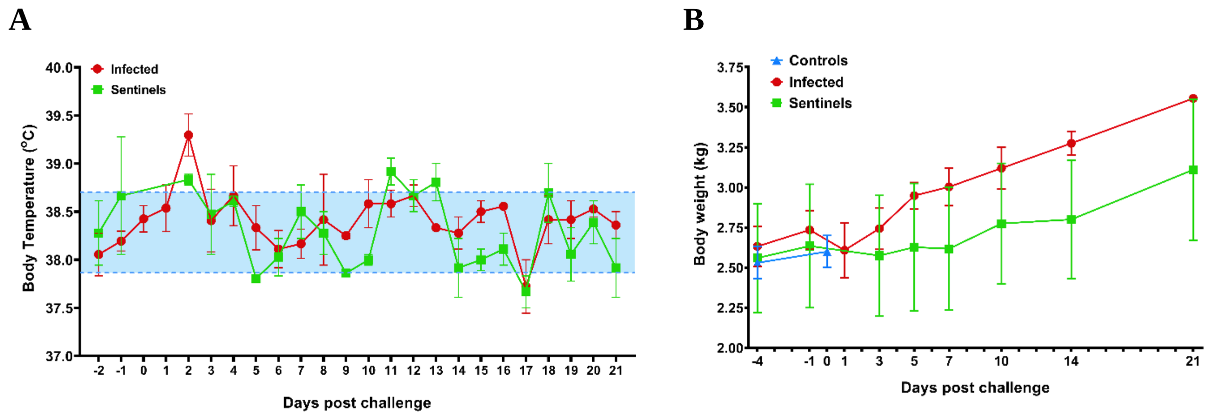


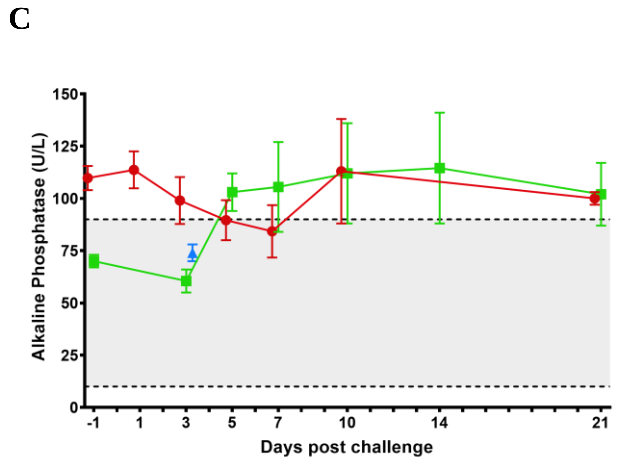


**Figure S1. Clinical parameters.** A) Body temperatures were recorded daily over the course of the study. B) Body weight of each cat was recorded throughout the study. C) Serum biochemistry and electrolyte analysis on 14 blood components was evaluated over the course of the study; elevated Alkaline Phosphatase levels are shown. Treatment group averages and standard deviations are shown in each panel. Shaded areas indicate normal range limits.

**A**

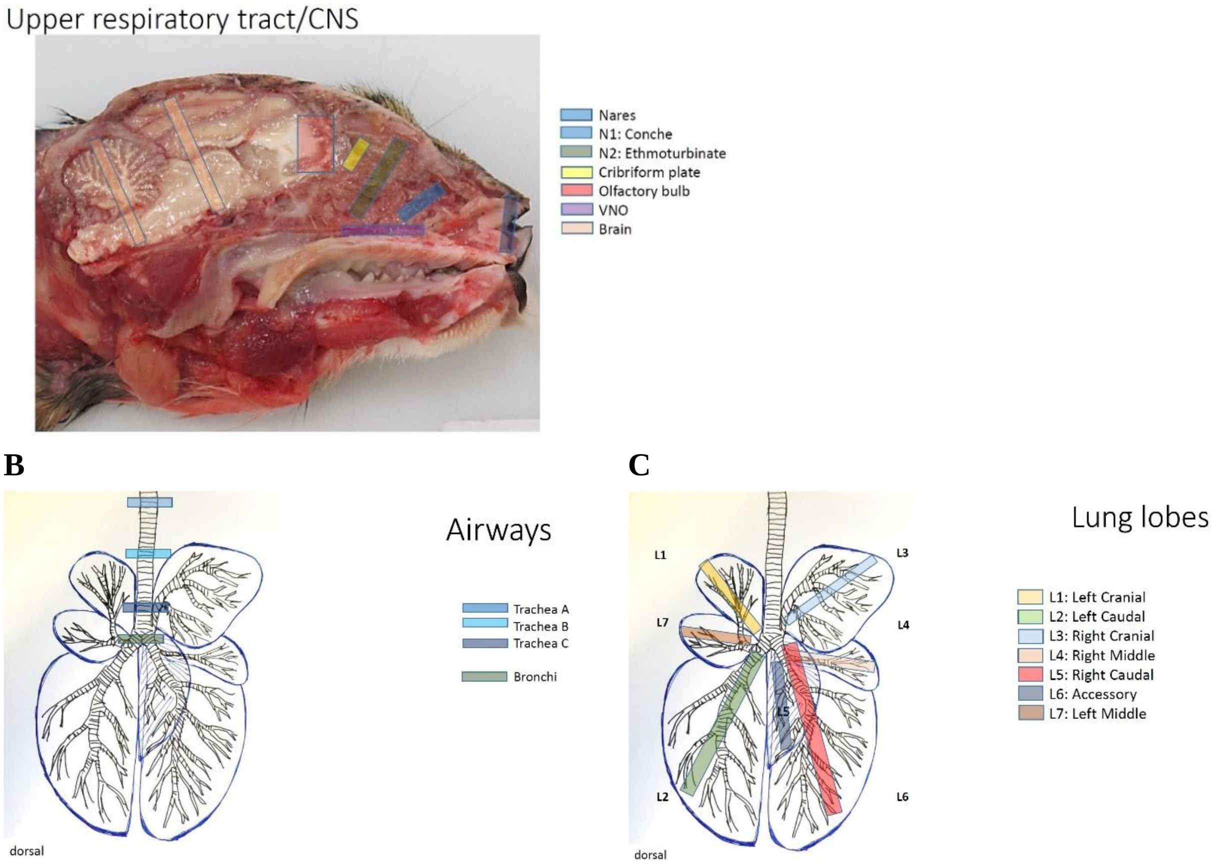


**Figure S2. Schematic of upper and lower respiratory tract tissues collected at necropsy.** Cross-sections of the central nervous system (CNS), nasal cavity, airways and lung lobes are indicated.

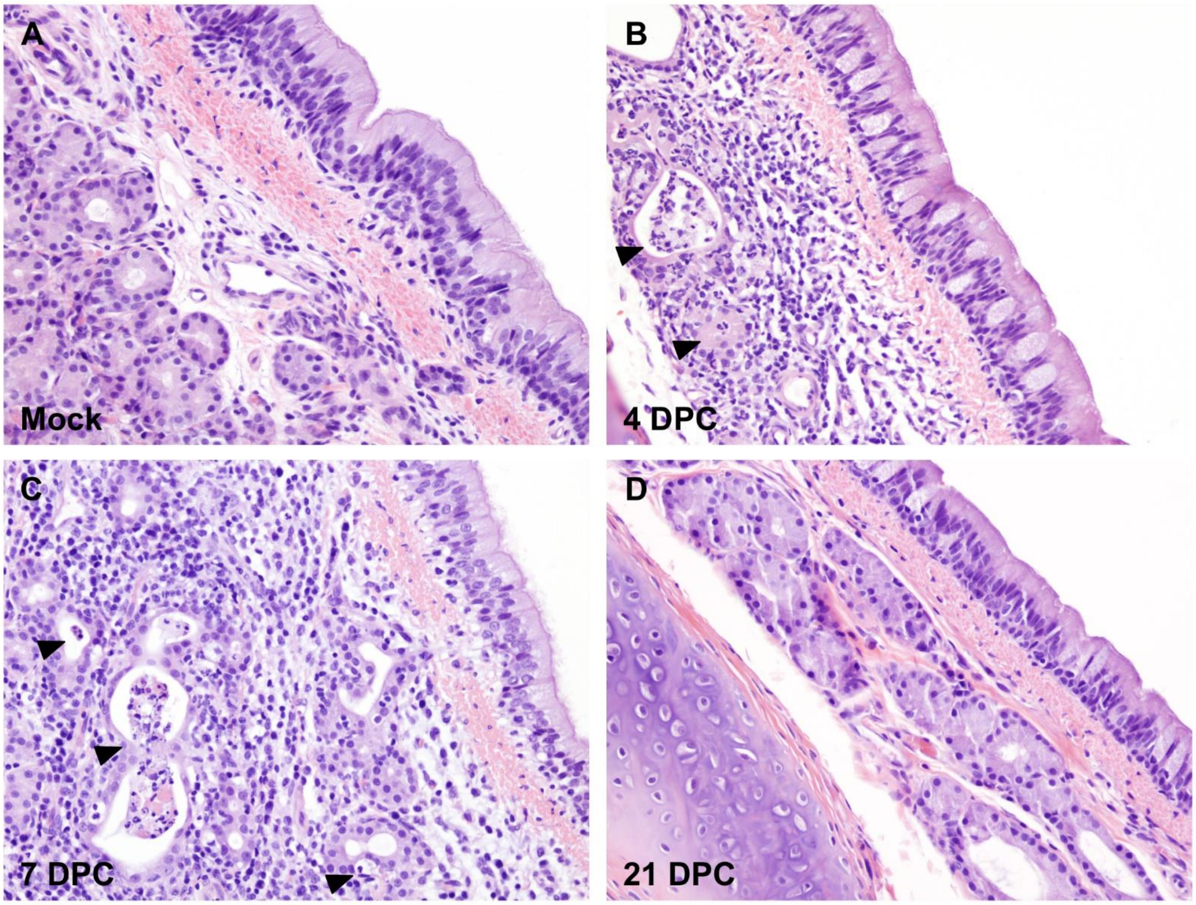


**Figure S3. Histopathology of trachea.** Histological findings in the trachea of mock (A) and SARS-CoV-2 experimentally infected (B to D) cats. At 4 days post-challenge (DPC; B) and 7 DPC (C), multifocal submucosal glands and ducts are distended (ectatic), filled with a small amount of necrotic debris, and lined by an attenuated epithelium (B and C arrowheads). The circumjacent interstitium is infiltrated by mild to moderate numbers of lymphocytes, histiocytes and scattered neutrophils that occasionally transmigrate through the lining epithelium. Inflammatory changes progress from minimal/mild to moderate between 4 and 7 DPC. No histologic changes are noted at 21 DPC (D). H&E. Total magnification: 200X

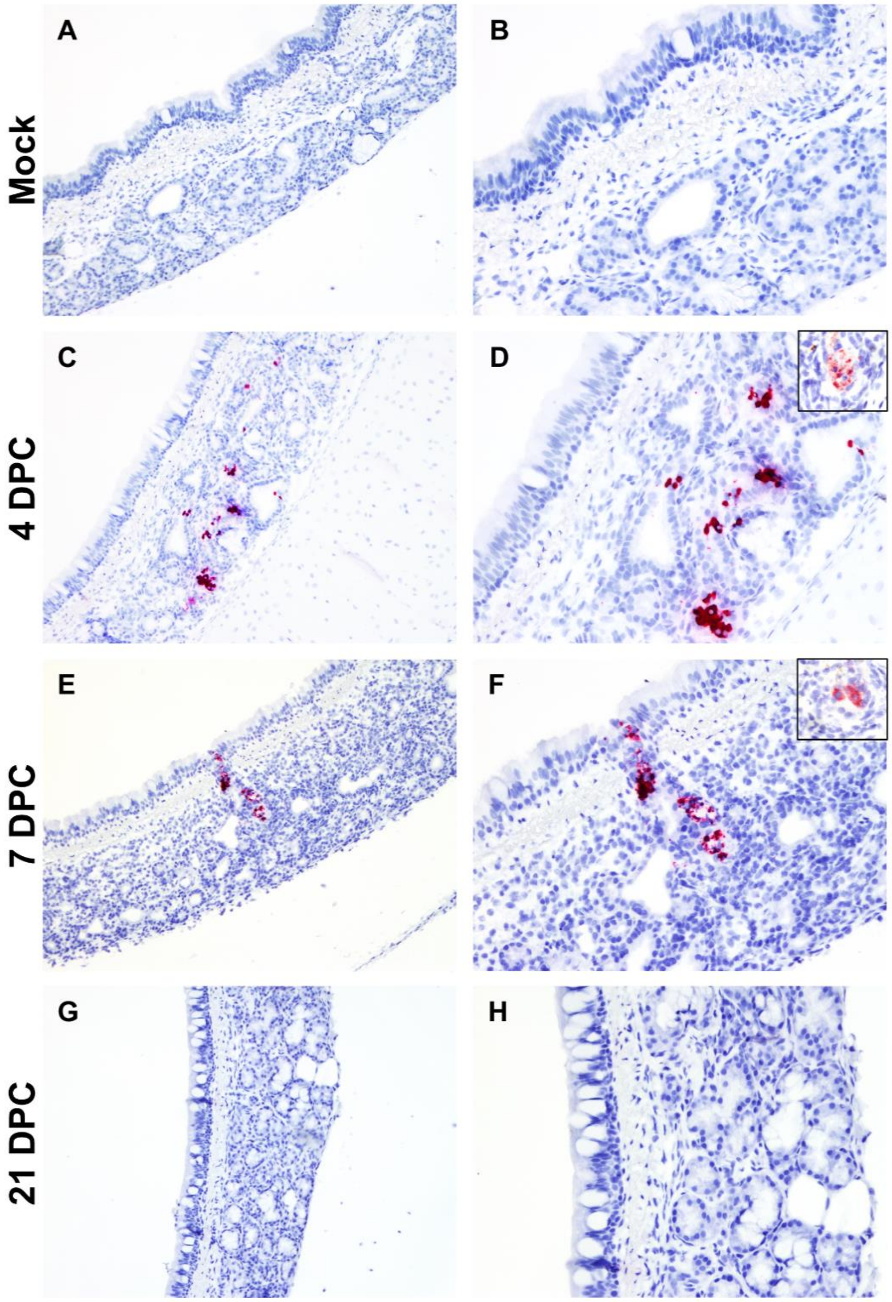


**Figure S4. SARS-CoV-2 RNA and antigen detection in trachea.** SARS-CoV-2 tropism in the trachea of mock (A and B) and experimentally infected (C-H) cats as determined by S-specific RNAscope® *in situ* hybridization (Fast Red) and anti-N-specific immunohistochemistry (IHC Fast Red). Glandular and ductular epithelial cells of multifocal, scattered submucosal glands are positive for viral RNA at 4 days post-challenge (DPC; C and D), and 7 days DPC (E and F). Similarly, few scattered glandular epithelial cells are positive for SARS-CoV-2 N antigen by IHC (D and F, insets). No viral RNA or antigen is detected at 21 DPC (G and H). Total magnification: 100X (A, C, E and G) and 200X (B, D, F and H).

10
